# Spatial Intra- and Intercellular Alignment of Respiratory Cilia and its Relation to Function

**DOI:** 10.1101/735332

**Authors:** Martin Schneiter, Sebastian Halm, Adolfo Odriozola, Helga Mogel, Jaroslav Rička, Michael H. Stoffel, Benoît Zuber, Martin Frenz, Stefan A. Tschanz

## Abstract

Ciliary alignment is considered necessary to establish respiratory tract mucociliary clearance, and disorientation is often associated with primary ciliary dyskinesia. Therefore, there is an urgent need for a detailed analysis of ciliary orientation (CO). We used volume electron microscopy to examine CO relative to the tracheal long axis (TLA) by measuring the inter- and intracellular basal body orientation (BBO) and axonemal orientation (AO), which are considered to coincide, both equivalently indicating the effective stroke direction. Our results, however, reveal that only the mean BBO is aligned with the TLA, whereas the AO determines the effective stroke direction as well as the mucociliary transport direction. Furthermore, we show that even if the mean CO is conserved across cell boundaries, a considerable gradient in CO exists within individual cells, which we suspect to be crucial for the emergence of coordinated ciliary activity. Our findings provide new quantitative insight into CO and correlate this new structural information with mucociliary function.

## Introduction

The epithelium of the tracheobronchial tree constitutes a self-cleaning surface, which mainly consists of secretory goblet cells and ciliated cells. Dust and infectious particles enter the respiratory tract when breathing. Due to adhesion, these particles get entrapped by the mucus layer lining the inner surface of the airways. Coordinated movements of billions of subjacent cilia then propel the overlying mucous carpet towards the larynx, thereby cleaning the lungs from inhaled substances.

Cilia are hair-like protrusions of the cell membrane with a particular cytoskeletal scaffold called axoneme. The ciliary axoneme consists of an evolutionary preserved 9+2 microtubular structure: a central pair is surrounded by nine circularly aligned microtubule doublets. The microtubules arise from the basal bodies, which are anchored to the cytoskeleton and serve as nucleation sites for the growth of axonemal microtubules. The chiral structure of the axoneme as well as of the basal foot appendage allows to unequivocally determine ciliary orientation (CO).

The physical orientational alignment of motile cilia along the proximal-distal axis of the airways is a prerequisite for concerted directional ciliary movement and the generation of directed fluid flow (e.g. Vladar et al. (2015); Guirao and Joanny (2007)). Studies attempting to quantify the orientation of respiratory cilia usually determined the direction of the assumed ciliary beating plane, or more specifically, the direction of the ‘effective stroke’ (or ‘power stroke’). According to the literature, the effective stroke direction is unambiguously defined, and can be inferred from two chiral ciliary structures: 1) the ciliary beating plane is assumed to be perpendicular to the central pair of microtubules, and in particular, the effective stroke is assumed to be directed from doublet 1 towards the gap between doublet 5 and 6 Satir and Christensen (2007); Satir et al. (2014). 2) The direction indicated by the tip of the basal foot appendage is considered to point into the direction of the effective stroke (e.g. Vladar et al. (2015); Gibbons (1961); Satir and Dirksen (1985)), which is sometimes also supposed to indicate the direction of fluid flow (e.g. Marshall and Kintner (2008); Chien et al. (2013); Satir and Dirksen (1985)). Moreover, the ciliary effective stroke direction is commonly considered to coincide with the direction of fluid flow. Consequently, many authors act on the assumption of a coincidence between the effective stroke direction inferred from the axonemal orientation, the effective stroke direction inferred from the basal foot orientation and the direction of fluid flow.

Ciliary disorientation has been proposed and discussed as a variant of primary ciliary dyskinesia (PCD) Rutman et al. (1993); Rayner et al. (1996); Rutland and De Iongh (1993); De Iongh and Rutland (1989). These studies particularly reported on patients showing all the clinical symptoms of PCD including abnormal or absent mucociliary clearance, while displaying normal ciliary beat frequency, normal ciliary beat pattern as well as normal ciliary ultrastructure, but exhibiting a disorganized CO as the only diagnostic finding indicating a disorder. In a more recent comprehensive study, the secondary nature of ciliary disorientation has been demonstrated Jorissen and Willems (2004). Therefore, as suggested in Marshall and Kintner (2008), the most conservative conclusion is that the importance of CO as an ultrastructural defect in PCD subjects remains to be fully explored. It is widely accepted that diagnosed disorganized ciliary orientation may either be the cause for mucociliary dysfunction, or a consequence of mucociliary anomalies. Ultimately, ciliary disorientation seems to be correlated with abnormal mucociliary clearance. It has very recently been shown that pathogenic variants in growth arrest-specific protein 2-like 2 (GAS2L2) lead to a clinical PCD-phenotype exhibiting normal axoneme structure, but uncoordinated hyperkinetic ciliary activity and impaired CO, which results in inefficient mucociliary clearance Bustamante-Marin et al. (2019). These recent findings thus underline the relevance of proper CO for efficient mucociliary clearance. Nevertheless, ciliary disorientation in the absence of other ultrastructural defects is presently not approved as a criterion for a positive PCD diagnosis, neither by the American Thoracic Society Shapiro et al. (2018), nor by the European Respiratory Society Lucas et al. (2017).

Current research on CO is typically focused on the identification of the governing mechanisms establishing initial polarization cues and guiding the common orientation of cilia during development Luo et al. (2017); Herawati et al. (2016); Chien et al. (2013); Guirao et al. (2010); Vladar et al. (2012); Werner et al. (2011); Mitchell et al. (2007). Planar cell polarity (PCP) proteins were shown to be asymmetrically localized in various organs and species, prior to the onset of ciliary beating, and are therefore suspected to establish initial directionality Vladar et al. (2012); Baumann (2010); Wallingford (2010). Basal bodies were particularly found to be highly disorganized (in terms of position and/or orientation) when docking at the apical cellular surface in different ciliated epithelial tissues of various species Spassky and Meunier (2017); Herawati et al. (2016); Guirao et al. (2010). Generally, it is considered that interactions of the actin and microtubular cytoskeletal network with PCP components set a polar bias Werner and Mitchell (2012); Mitchell et al. (2007). The actin and microtubular network is furthermore considered to connect, reorient and (spatially) realign the basal feet Vladar et al. (2012); Werner and Mitchell (2012); Werner et al. (2011). The onset of ciliary beating is then thought to refine CO by a positive feedback mechanism: hydrodynamic forces created by the collectively generated fluid flow refines the orientation of roughly oriented young cilia in a common direction, which in turn amplifies fluid flow Guirao et al. (2010); Mitchell et al. (2007).

The motivation for the present paper originates from the formulation and analysis of an oversimplified pluricellular epithelium model Schneiter et al. (2019); Ricka (2010), which represents an array of locally interacting ciliated cells able to self-organize towards a self-cleaning virtual epithelium. When ‘constructing’ such a virtual epithelium, modelers need access to dedicated quantitative morphological information. Despite the available large body of literature concerning CO during development, the effect of cellular boundaries and of the spatial ciliary alignment on the orientation of the ciliary beating plane remains poorly characterized. Therefore, we conjectured different conceivable geometries, each representing a specific inter-cellular organization of the ciliary beating plane. Consequently, we were left with an unresolved question: does the mean orientation of cilia vary between neighboring cells (possibly even according to a specific inter-cellular spatial pattern)? Or, is CO consistent beyond cell boundaries? In order to reach the ultimate goal of the accurate understanding of mucociliary phenomena, numerical models require access to this kind of dedicated morphological information, particularly with respect to the spatial organization of CO. The present work intends to fill the gap of information regarding the spatial organization of CO and to relate the structural features of CO to functional behavior such as mucociliary transport.

## Results

### Ciliary Orientation with Respect to the Tracheal Long Axis

An overview of all determined angular values for the basal body orientation (n = 1661) and the axonemal orientation (n = 1998), is provided in Table 1 and Table 2, respectively. The data was divided into four groups, which correspond to the four stages of the sample preparation (see Fig.1). Consequently, Table 1 and Table 2, which list the values for the circular mean and standard deviation within each group, are organized as follows: the values derived from the same *field of view*, i.e. from the same image stack, are listed in the bottom row (row 4). The distribution of the measured angles *ω* (BBO) and Θ (AO) within each field of view are additionally illustrated in terms of radial histograms in Table 1 – Figure Supplement 1 and Table 2 – Figure Supplement 2, respectively. For the next higher level (row 3), the values derived from the *same block*, originating either from CT or LA, were summarized. The angles derived from the two blocks, which correspond to the *same animal* were summarized in row 2. Finally, the top row (row 1) corresponds to the largest group ofdata, for which all the values derived from all four animals (≙ 8 blocks and 16 field of views) were combined.

**Table 1:**
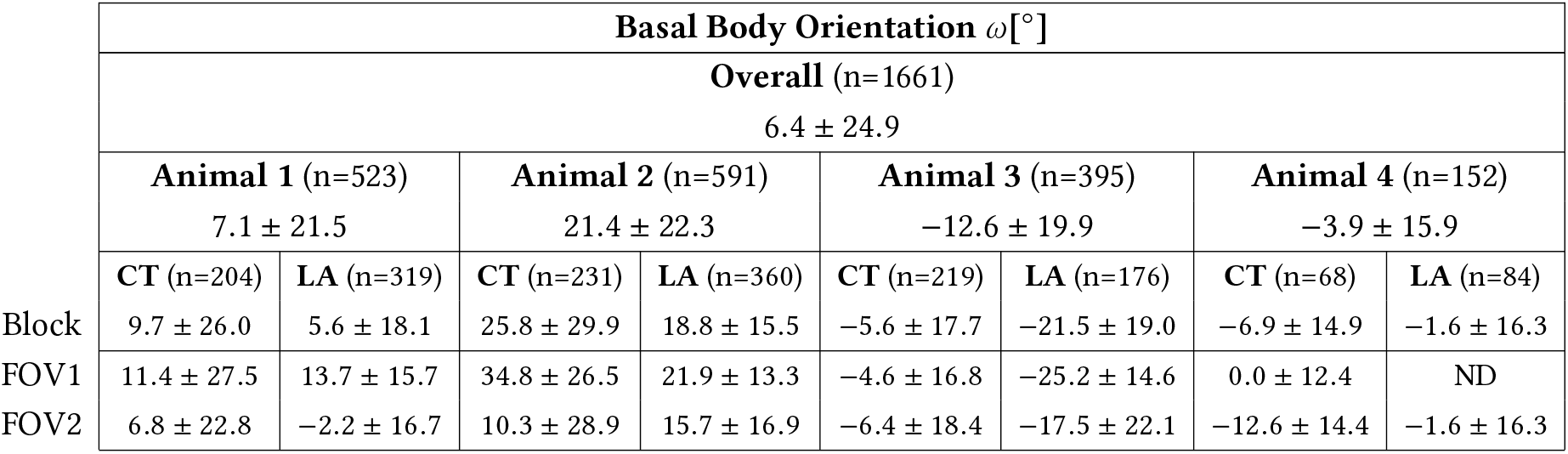
Overview of the measured basal body orientations (*Θ*). Each cell contains the circular mean and the circular standard deviation (values are listed in [°]) within the indicated group (rows from top to bottom: overall, per animal, per block sample and per field of view). The found basal body orientations are additionally graphically represented in Table 1 – Figure Supplement 1.

**Table 2:** Overview of the measured axonemal orientations (Θ). Each cell contains the circular mean and the circular standard deviation (values are listed in [°]) within the indicated group (rows from top to bottom: overall, per animal, per block sample and per field of view). The found axonemal orientations are additionally graphically represented in Table 2 – Figure Supplement 1.

| Axonemal Orientation $\Theta[^\circ]$ | | | | | | | |
| --- | --- | --- | --- | --- | --- | --- | --- |
| Overall (n=1998) |  |  |  |  |  |  |  |
| 19.4 ± 23.2 | | | | | | | |
| Animal 1 (n=386) |  | Animal 2 (n=370) |  | Animal 3 (n=670) |  | Animal 4 (n=572) |  |
| 24.9 ± 18.0 | | 41.1 ± 25.9 | | 9.6 ± 21.6 | | 14.0 ± 15.5 | |
| CT (n=133) | LA (n=253) | CT (n=127) | LA (n=243) | CT (n=407) | LA (n=263) | CT (n=255) | LA (n=317) |
| Block 25.2 ± 22.7 | 24.8 ± 15.1 | 60.3 ± 34.1 | 32.7 ± 13.5 | 11.3 ± 16.5 | 6.6 ± 27.6 | 11.1 ± 15.6 | 16.3 ± 15.1 |
| FOV1 29.5 ± 14.7 | 29.9 ± 15.1 | 73.3 ± 26.7 | 34.0 ± 11.7 | 13.6 ± 17.2 | 5.3 ± 22.7 | 19.9 ± 10.2 | 14.7 ± 15.5 |
| FOV2 17.9 ± 30.1 | 20.6 ± 13.7 | 41.1 ± 34.5 | 30.8 ± 15.6 | 9.0 ± 15.5 | 8.8 ± 33.7 | 3.0 ± 15.4 | 17.1 ± 14.8 |

**Figure 1:**
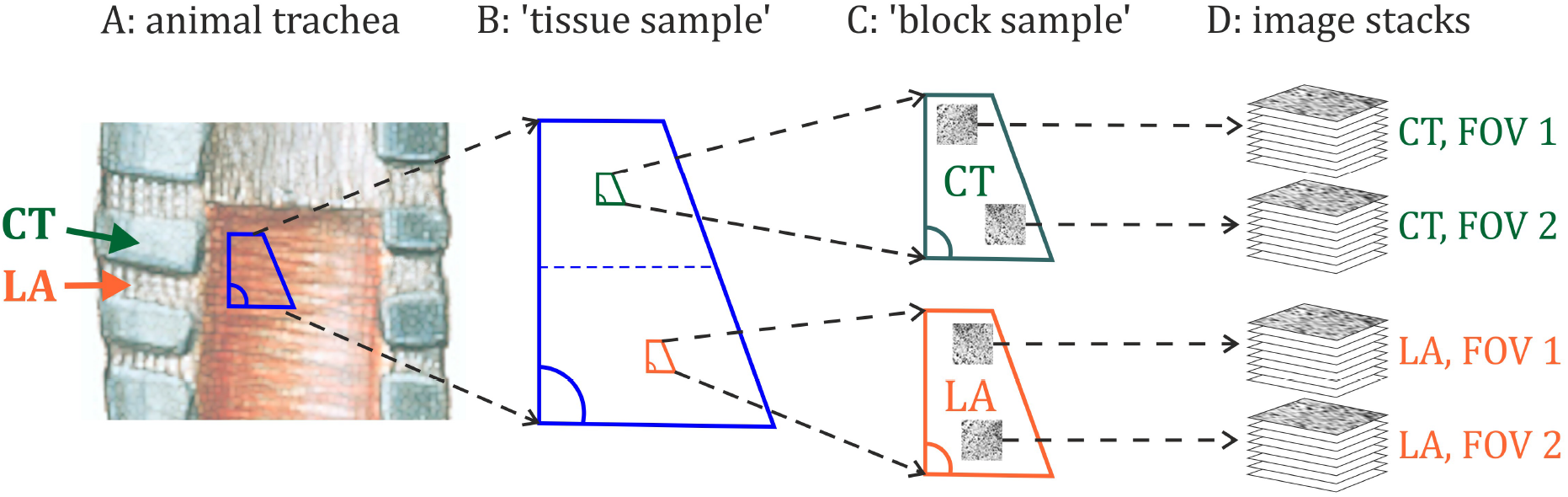
Four procedural substeps (A,B,C,D) of the sample preparation are illustrated. A: one large trapezoidal mucous membrane sample (≈ 3cm×2cm) was excised from each trachea. As indicated, each large trapezoid, which is referred to as the ‘tissue sample’ in the following, comprises epithelial tissue overlying tracheal cartilages (CT) as well as annular ligaments (LA). B: from each large fixed trapezoidal epithelium sample, two smaller (≈ 3mm×2mm) trapezoids were then excised: one being associated with CT-tissue and one with LA-tissue. These smaller trapezoidal samples are referred to as ‘block samples’ in the following. C: after processing the tissue according to the protocol for serial block face scanning electron microscopy, two regions of interest on each of the block samples were imaged. D: consequently, from each trachea four image stacks (2×CT, 2×LA) were produced. As indicated, each image stack can be regarded as a (three-dimensional) field of view (FOV).

Overall, we found the mean basal body orientation to be 〈*ω*〉_*c*_ = 6.4° and the mean axonemal orientation to be 〈Θ〉_*c*_ = 19.4°. It should be noted that these circular mean values were derived by equally weighting each cilium. In consideration of the sample preparation, however, one would receive more meaningful estimates 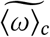 and 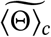 of the overall mean directions by weighting the mean values derived from each of the small trapezoidal block sample (summarizing therefore two fields of view derived from the same block, which corresponds to the third row in Table 1 and Table 2) according to their corresponding standard error (SE). The statistical SE of the mean direction determined for each sample block is considerably lower than the SE introduced by the practical approach, i.e. by the excision of the larger trapezoidal tissue samples and by the positioning of the smaller trapezoids.

Denoting the block-wise derived mean values as *γ*_1_,…, *γ*_8_, the sampling-corrected estimate and its SE can be calculated according to Eq.8 and Eq.9 (stated in Materials & Methods), respectively. The ‘sampling-corrected’ estimate for the mean basal body orientation, and its SE, finally amounts to: 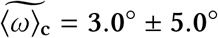. The ‘sampling-corrected’ estimate for the mean axonemal orientation, and its SE, amounts to: 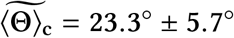.

The mean difference between the axonemal and the basal body orientation, which is denoted as 〈Δ〉_*c*_ = 〈Θ − *ω*〉_*c*_ was estimated as follows. One should note that Δ represents a relative measure and, thus, is not affected by errors introduced by the preservation of the tracheal long axis (TLA). Consequently, each sample block *i* delivers a certain value for the relative difference between the axonemal orientation and the basal body orientation, which we denote as Δ_*i*_. The associated SE *σ*_Δ_*i*__ was calculated according to: 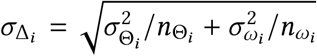, where *n*_Θ_*i*__ and *n_ω_i__* denote the number of measurements for the AO and the BBO performed in the sample block *i*, respectively. *σ*_Θ_*i*__ and σ_*ω*i_ denote the respective standard deviations. As the difference Δ represents an angular observable, i.e. an intermediate angle, each Δ_*i*_ was transformed into the unit vector 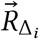, which was subsequently weighted by its inverse SE. The mean difference 〈Δ〉_*c*_ is finally given by the direction of the weighted mean resultant vector 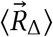:

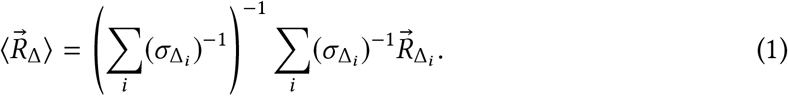

The SE of the mean difference was estimated by the pooled SE. This approach yields the following estimate for the mean difference between the axonemal and the basal body orientation: 〈Δ〉_*c*_ = 19.1° ± 2.1°.

In conclusion, we found that the tip of the basal foot and the arrow drawn perpendicular to the central pair orientation, encloses an angle of 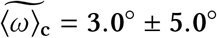 and 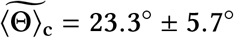 with the tracheal long axis pointing towards the larynx, respectively. While the laryngeal-directed TLA is contained in the *σ*-confidence interval for the mean BBO, the mean AO differs with a high significance (at the 4*σ*-level). The difference between the two highly significant distinct orientational observables amounts to 〈Δ〉_*c*_ = 19.1° ± 2.1°. It has to be mentioned that neither the mean AO nor the mean BBO showed a statistically significant difference between samples collected from above the cartilage rings (CT) and from above the tracheal ligaments (LA).

### Characteristic Directions of Mucociliary Function

In order to discuss the meaning of the two structurally derived orientational measures, i.e. the mean AO and the mean BBO, with respect to mucociliary function, we relate the morphological data to functional observables characterizing the oscillations of the mucous surface. The functional data, presented in the following, was derived from previous experimental work on bovine trachea explants Burn (2009). As the experimental procedure as well as imaging setup based on high-speed reflection contrast microscopy was previously described in Ryser et al. (2007) with additional technical information in Burn (2009), the principles are only briefly outlined in the Materials & Methods section.

Fig.2 summarizes the set of functional parameters derived from 56 measurements and 14 tracheas (1-7 fields of view were captured for one second on each trachea). The top row in Fig.2 emphasizes that each of the derived transport and wave propagation velocities represent a vector having a certain magnitude as well as a certain direction. The second row in Fig.2 shows the compilation of the corresponding radial histograms, which neglect the magnitudes and, therefore, represent the distribution of the transport direction and the wave propagation direction.

**Figure 2:**
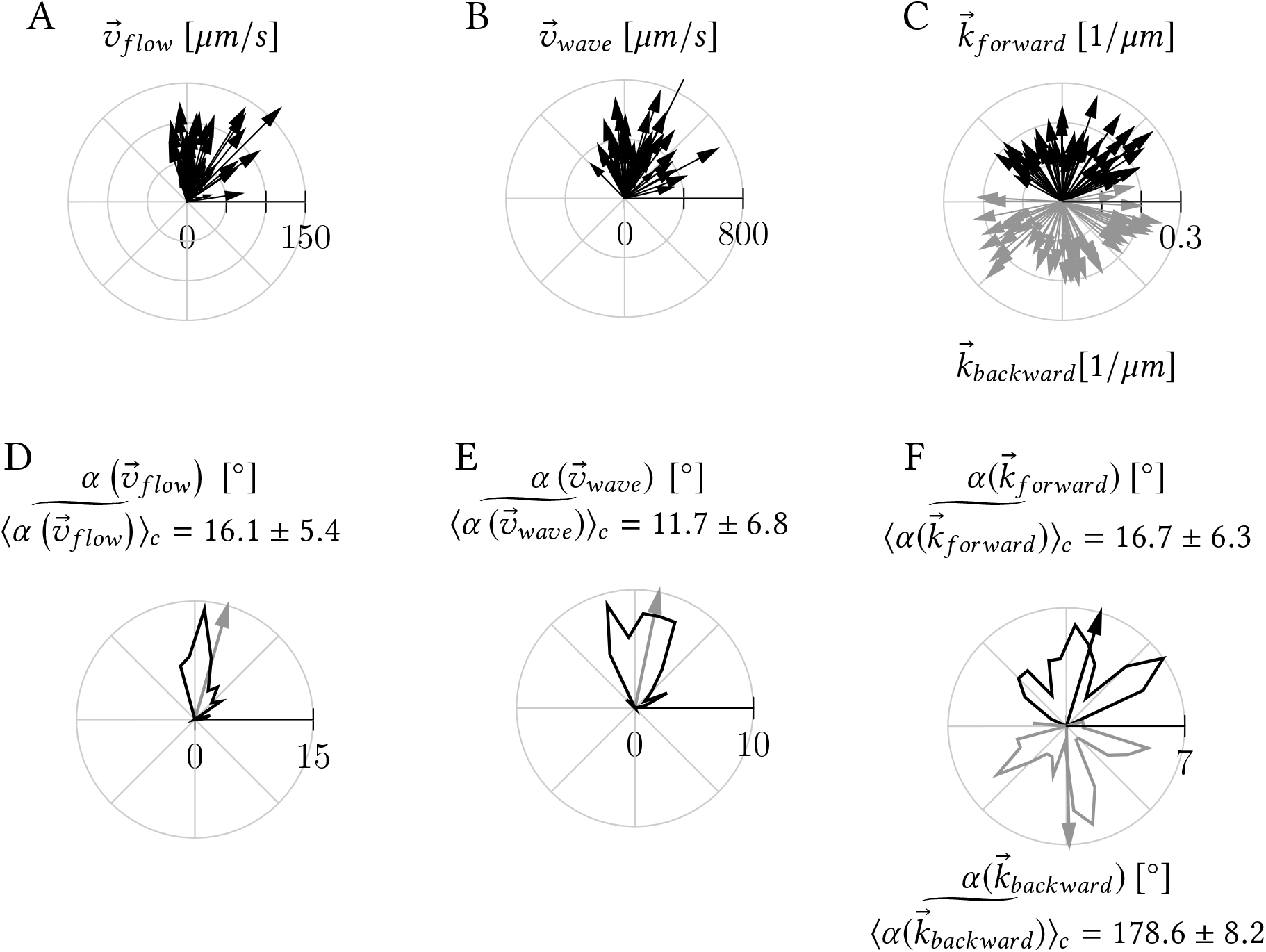
A: mean tracer transport velocities representing the mucociliary flow velocities 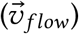. B: distribution of the propagation velocity of the ‘mean wave’ 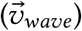. C: mean wave vectors representing the ‘mean harmonic plane wave’ propagating into the pharyngeal direction (black vectors) and the ‘mean harmonic plane wave’ propagating into the opposite direction (gray vectors). D-F: radial histograms generated by the extraction of the corresponding propagation directions displayed in the first row (A-C). The arrows represent the respective sampling corrected circular mean values.

### Differences in Ciliary Orientation Between Neighboring Cells

In order to uncover whether the mean CO changes between neighboring cells, we examined changes in CO between cells for which it was possible to determine the orientation of a minimum of 25 cilia (AO- or BBO-values). Moreover, only pairs of cells derived from the same field of view were compared with each other.

The histogram shown in Fig.3 provides an overview of how much intracellular mean directions deviate between neighboring cells. In total, the distribution of 35 absolute differences between intracellular means as inferred from the AO and 17 absolute differences between intracellular means as inferred from the BBO is illustrated. It is evident that intracellular mean orientations differ only slightly between cells: the median of the absolute difference of the AO and the BBO amounts to 4.9°. This value reflects a remarkable highly organized CO on a pluricellular level.

**Figure 3:**
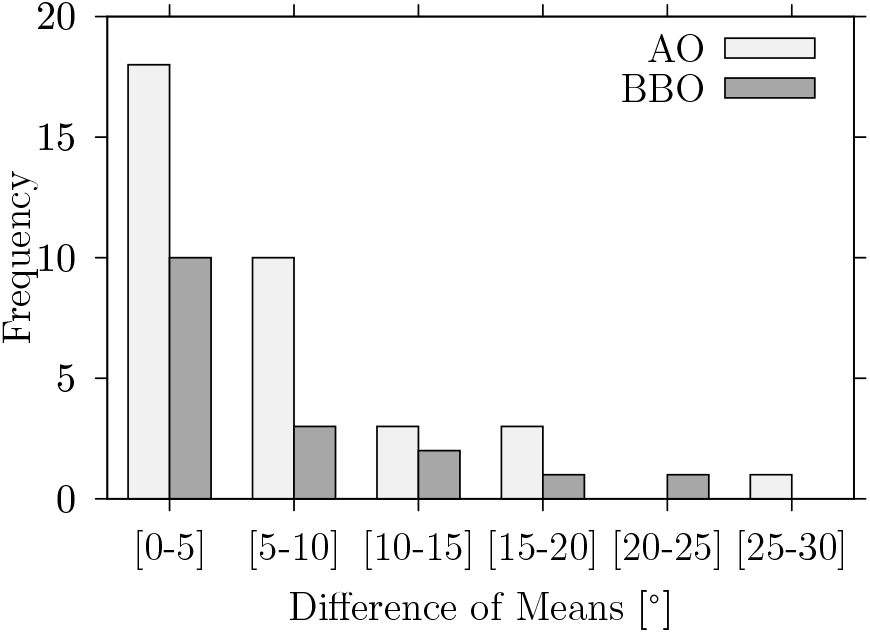
The histogram shows the distribution of the absolute difference between intracellular means in cilia orientation (AO- and BBO inferred values). All values correspond to pairs of cells, which correspond to the same field of view. Only cells with *n* ≥ 25 were considered.

In order to test whether pairs of cells share the same mean direction, we calculated Welch’s *t*-intervals with a significance level of *α* = 1%. The resulting confidence intervals for the difference between intracellular means are illustrated in Fig.4. These confidence intervals reflect an estimate, how much the difference between intracellular mean values is expected to vary, based on the collected data samples and the assumption that the data are random samples from two populations with a coinciding mean. Confidence intervals not crossing the red line indicate that the respective differences of intracellular means significantly differ from each other and, thus, provide the same information as significant p-values (at *α* = 1%) derived from independent two-sample *t*-tests with unequal variances.

**Figure 4:**
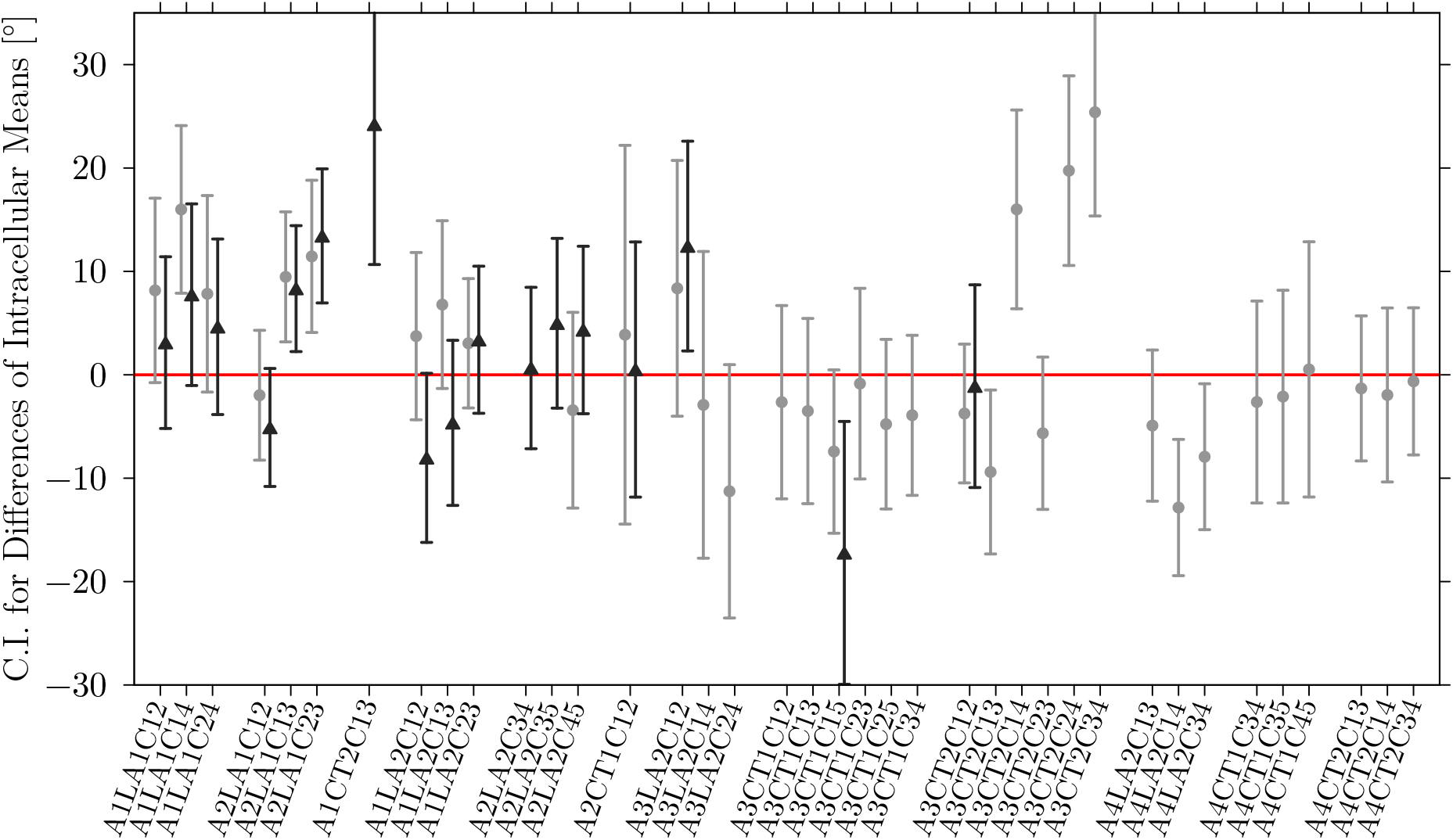
The graph shows 99%-Welch’s *t*-confidence intervals for the difference between intracellular means. Light gray and dark gray intervals correspond to AO and BBO values, respectively. The dots and triangles indicate the difference between intracellular means in AO and BBO, respectively. The x-tick labels specify which pair of cells are being compared: for instance, the label ‘A1LA2C23’ denotes the difference of intracellular means between cell 2 and cell 3 (‘C23’) determined on the second image stack, which was derived from ligamentum annulare (‘LA2’) from animal 1 (‘A1’).

Fig.4 shows that about three-fourths of all confidence intervals cross the red line indicating that the corresponding observed differences between intracellular mean power stroke directions do not differ significantly.

When interpreting the partially significant difference between intracellular means, the following conditions must be taken into account: 1) In a few cells the determined values for *ω* and Θ deviate considerably from a normal distribution. Thus, the assumption of intracellular normally distributed values was not always met. 2) The sample size, i.e. the number of COs we were able to determine per cell, varied between *n* = 25 and *n* = 84. Highly significant differences of means correspond to pairs of cells comprising at least one cell, for which only a low number of COs was available. On the other hand, mean directions corresponding to pairs of cells with larger sample sizes show almost coinciding mean effective stroke directions, even though statistical power grows with an increasing sample size. 3) As we will see, the apparent contradiction from the last point gets resolved by the fact that our sampling was not random (biased representation of the population), which is caused by spatially non-isotropically distributed data being subject to spatial correlations.

In order to complement the pairwise comparisons between intracellular means, we used a global descriptive approach to analyse the impact of cell boundaries on CO. This additional approach is based on the comparison of the orientational discrepancy in-between pairs of cilia picked from the same cell with the discrepancy in-between pairs of cilia picked from distinct cells. Therefore, two sets of angular deviations (in BBO and AO) within each possible pairwise combination of cilia were constructed: 1) the set of angular deviations within each possible intracellular combination of cilia pairs, which are referred to as *IntraBBO* and *IntraAO*, and 2) the set of angular deviations within all possible combinations of intercellularly drawn pairs of cilia, which are referred to as *InterBBO* and *InterAO*. The construction of these combinatorial sets is formally denoted in Eq.2 and Eq.3,

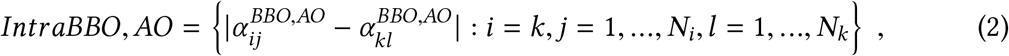

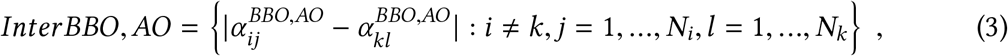

where *BBO, AO* indicates that the sets were constructed with respect to both orientational observables. Theindicesi andk specifythe cellandthe indices *j* and *l* iterate over all cilia of cell *i* and *k*, respectively. The sets *IntraBBO, IntraAO* and *InterBBO, InterAO* were constructed for each field of view separately. The boxplots shown in Fig.5 illustrate the distribution of intracellularly and intercellularly drawn pairs of cilia collected over all fields of view (> 5 · 10^5^ values). The boxplots indicate a slight tendency to larger deviations for pairs of cilia drawn from distinct cells (for both orientational measures): the median deviation in IntraBBO and InterBBO amounts to 15.2° and 15.8°, respectively, and in IntraAO and InterAO to 13.8° and 15.4°, respectively. Considering the bumpy apical cellular surface and that cilia drawn from distinct cells are located considerably further apart than cilia drawn from the same cell, the overall directional similarity between intra- and intercellular deviations in CO indicates that the mean CO does not change between neighboring cells.

**Figure 5:**
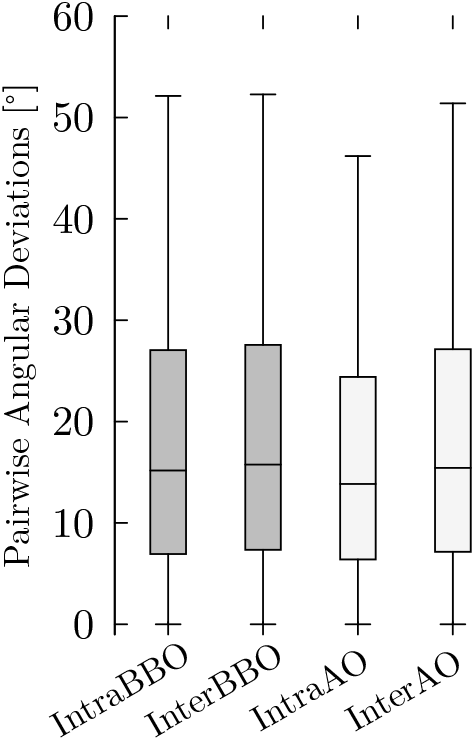
Boxplots illustrating the distribution of angular deviations within pairs of cilia (from the left to the right): drawn from the same cell in terms of the BBO, drawn from different cells in terms of the BBO, drawn from the same cell in terms of AO and as well as from different cells in terms of AO. The whiskers indicate the range covered by 95% of all data, outliers were suppressed.

### Spatial Correlations in CO

In consideration of Tobler’s first law of geography Tobler (1970): ‘*everything is related to everything else, but near things are more related than distant things*’, we used geostatistical methods in order to investigate the spatial dependence of our spatially irregularly distributed data.

### Radial Variogram

In order to measure spatial similarity (or rather dissimilarity in our case) we made use of the so-called empirical madogram, which is a first order version of the empirical variogram Legendre and Legendre (1998); Mathur (2015). We used a radial (one-dimensional) version, denoted in the following as (|Δ*α*|)(*r* ± *δ_r_*), which provides a measure of how much the orientation of two cilia (*α_i_* and *α_j_*) separated by a certain distance deviate from each other and which is defined as:

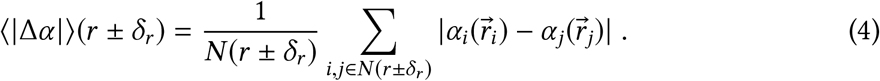

To put it simply, (|Δ*α*|)(*r* ± *δ_r_*) denotes the average angular deviation between pairs of cilia located at 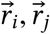, which fall into a certain distance class *r′* ± *δ_r_*, i.e. 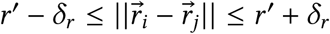. *N* (*r* ± *δ_r_*) denotes the number of all pairs of cilia located in respective distance classes. In practice, (|Δ*α*|)(*r* ± *δ_r_*) is constructed as follows: for *N* cilia all 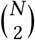 possible pairs are built and subsequently, the associated distances as well as the angular deviations are determined. After choosing a certain distance class width (2*δ*), the average angular deviation in each distance class is calculated. Note that the distance between two cilia located at 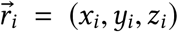 and 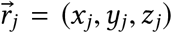, were calculated such that it approximately represents the ‘geodesic distance’ along the curved cellular surface.

### Directional Variogram

Besides the variogram version defined in Eq.4, we used a directional version of Eq.4 delivering two-dimensional variograms, denoted as (|Δ*α*|)(Δ*x* ± *δ_x_*, Δ*y* ± *δ_y_*), which was calculated according to:

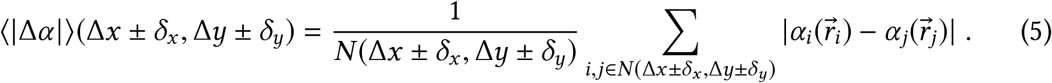

The directional, or two-dimensional, variogram was applied in its unsigned version (according to the formulation in Eq.5), as well as in its signed version, i.e. 〈Δ*α*〉_*c*_(Δ*x* ± *δ_x_*, Δ*y* ± *δ_y_*). The horizontal and vertical distances between two cilia located at 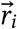 and 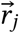, i.e. Δ*x_ij_* and Δ*y_ij_*, were determined after geodesically projecting the coordinates of the respective cilia. In simple terms, the three-dimensional curved cellular surface was flattened by an approximate geodesic mapping on a two-dimensional plane. Subsequently, the spacing between the projected coordinates, denoted by Δ*x* and Δ*y*, were determined.

### Direction of Intracellular Gradients in Cilia Orientation

In order to determine the direction of clockwise deflection between two cilia located at 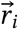 and 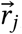, respectively, we introduced the angle *β*, which is defined as:

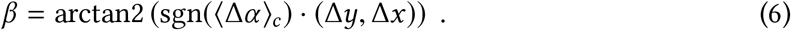

In Eq.6, sgn(*x*) denotes the sign function. Δ*α* represents the angular difference between cilium *i* and cilium *j*: 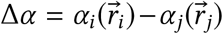. Finally, Δ*y* and Δ*x* are given by the projection of the relative distance vector 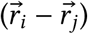 on the *y*- and *x*-axis, respectively.

A set of *N* spatially distributed cilia provides 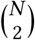 pairs of cilia, for which the directions of clockwise angular deviations were determined. The circular average over these directions yields the mean direction of clockwise deflections in cilia orientation, which is denoted by 〈*β*〉_*c*_.

### Intracellular Gradual Shift in Cilia Orientation

The variograms introduced in Eq.4 and Eq.5 are based on pairwise comparisons of CO with respect to the relative spatial positions of cilia. The introduced empirical variograms can be constructed from all possible pairs of cilia for which: 1) both cilia *i* and *j* are drawn from the same field of view (FOV), 2) both cilia *i* and *j* are drawn from the same cell, and 3) cilium *i* and cilium *j* are drawn from distinct neighboring cells. Thus, for each field of view (FOV), we generated 1) intra-FOV variograms, 2) intracellular variograms, as well as 3) intercellular variograms.

The various kinds of variograms, which were examined for all fields of view, are exemplarily illustrated in Fig.6A-C for axonemally derived orientations from sample ‘A4LA2’. A collection of additional representative variograms is provided in Figure 6 – Figure Supplement 1. The shape of the generated *radial variograms* revealed the following ubiquitous regularities across different samples:

1. *Orientational deviations increase with interciliary distances:* as shown in Fig.6A, the intracellular radial variogram (solid gray line) displays an increase of the orientational discrepancy among pairs of cilia from 10° to 25° with increasing relative distance. This means that closely aligned pairs of cilia are more similarly oriented than further distanced pairs. The pairwise angular deviation generally increased with an increasing interciliary distance. At interciliary distances above 4-6*μ*m a typical increase of 10-15° was observed.
2. *Intercellular (radial) variograms display no distinct behavior*: the dashed black curve in Fig.6A represents a typical intercellular radial variogram. Overall, the intercellular radial variograms indicate no intercellular spatial dependency in CO, as they display only little variation with no noticeable patterns.
3. *Cilia orientation de-correlates at interciliary distances of 4-6 μm:* the FOV-variograms (solid black line in Fig.6A) consistently show an initial increase, which is dominated by intracellular cilia pairs, with the interciliary distance and a correlation length of ≈ 4-6 *μ*m. At distances beyond the correlation length, the variogram is guided by the intercellular variogram.
4. *Maximum orientational discrepancy was found within cells:* It should be pointed out that intracellular cilia pairs separated by more than 4 *μ*m are considerably less similarly oriented than intercellular cilia pairs, which represents an unexpected finding.

The two-dimensional variograms provide directional information of pairwise orientational deviations. In general, the generated FOV-wide two-dimensional variograms did not show any distinct patterns across the available fields of view. Sample A4LA2 however suggests a ‘morphological wave-like pattern’ (see Fig.6B). Unfortunately, the available data sets can neither prove nor disprove such a pluricellular pattern. This would require to examine larger fields of view providing spatially isotropically distributed data.

As can be seen in Fig.6C, the signed intracellular variograms show evidence of an intra-cellular gradient in cilia orientation: CO gradually shifts in clockwise direction from the right to the left perpendicularly to the TLA (view on the epithelium of the ventral tracheal wall).

**Figure 6:**
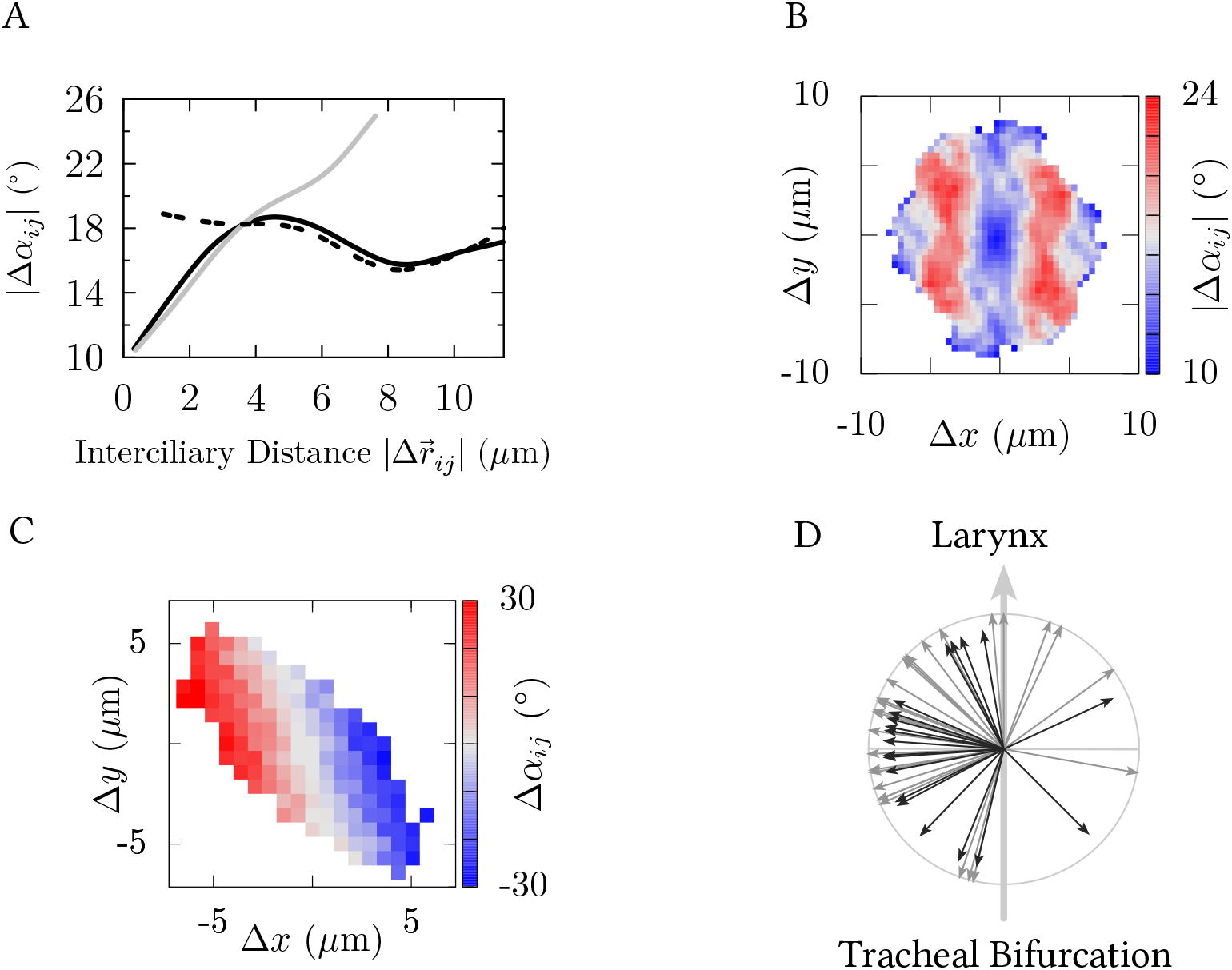
The panels A-C show the radial and directional variograms generated with the AO data derived from sample ‘A4LA2’. A: radial variograms (intra-FOV: solid black, intracellular: solid gray, intercellular: dashed black). B: absolute directional FOV-variogram. C: signed directional intracellular variogram. D: The arrows display the distribution of intracellular mean directions towards clockwise deflections in cilia orientationi, i.e. 〈*β*〉_*c*_. Gray arrows correspond to AO-derived directions (28 cells) and black arrows to BBO-derived directions (18 cells).

In order to prove that the alleged intracellular orientational gradient can not be caused by outliers, we used Moran’s I, which can be transformed to *z*-scores (e.g. Mathur (2015); Legendre and Legendre (1998)), to statistically test whether the direction in which the orientation of adjacent cilia (nearest neighbor) deviates in a random manner or not. The application of Moran’s I on signed intracellular directional variograms, i.e. 〈Δ*α*〉_*c*_ (Δ*x*, Δ*y*), delivered highly significant p-values. This confirms an intracellular gradual shift in cilia orientation, which we found ubiquitously in all the cells.

In approximately 90% of cells, for which we were able to measure at least 30 values for the AO or the BBO, cilia orientation gradually shifted in a clockwise direction when seen from the right to the left perpendicularly to the TLA, as shown in Fig.6D.

## Discussion and Conclusions

The use of a scanning electron microscope equipped with a 3view module allowed the two-fold measurement of the CO in terms of the axonemal orientation (AO), which was unambiguously derived from the central pair orientation (Fig.7), and in terms of the basal body orientation (BBO), which was determined by the direction indicated by the tip of the basal foot (Fig.8). The AO, as well as the BBO, were measured under preservation of the laryngeal direction along the tracheal long axis (TLA) and, in particular, the information of the cellular affiliation of each cilium.

**Figure 7:**
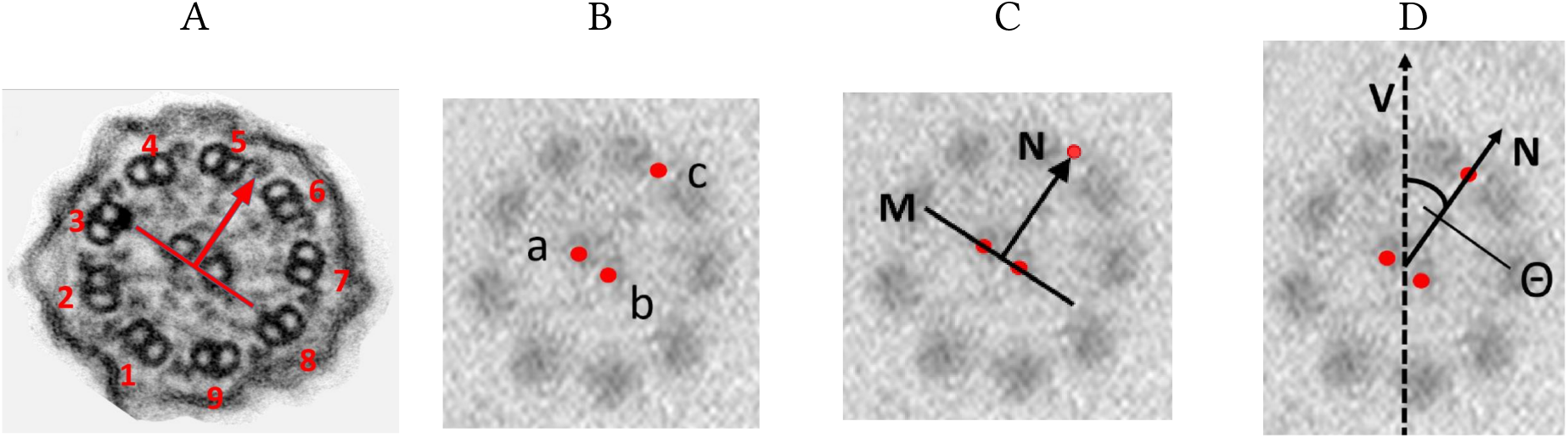
The panels A-D illustrate the measuring of the axonemal orientation. A: Transmission electron micrograph illustrating the assumed ciliary effective stroke direction being perpendicular to the line connecting the central pair and pointing from peripheral doublet 1 towards the gap between peripheral doublets 5 and 6. B: three points a,b and c labelling the coordinates of the central pair and the gap between peripheral doublets 5 and 6, respectively. C: line M connects the two central microtubules. The axonemal orientation is then indicated by the arrow N. D: Finally, Θ represents an unambiguous observable for the axonemal orientation with respect to the TLA, which coincides with the dashed vertical arrow V pointing towards the larynx.

**Figure 8:**
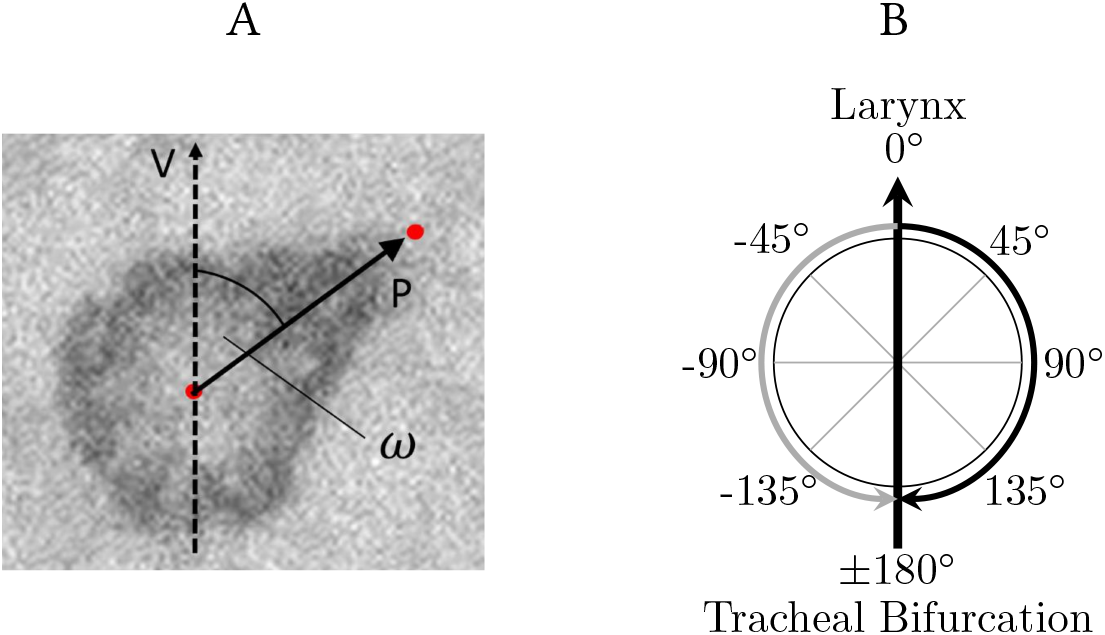
A: The basal body orientation was unambiguously defined by the basal foot orientation, which was measured by the angular observable *ω* enclosed by the arrow ‘P’ and the dashed arrow ‘V’. The arrow ‘P’ points from the center of the basal body to the tip of the basal foot and the dashed arrow ‘V’ coincides with the long axis of the trachea and points into laryngeal direction. B: the axonemal orientation as well as the basal body orientation were measured in terms of the angular deviation from the tracheal long axis pointing into laryngeal direction (vertical arrow).

We found that the tips of the basal feet point, in average, towards 3° ± 5°, when 0° indicates the TLA pointing in laryngeal direction. Positive values indicate a clockwise-directed deviation (note that the tracheal tissue samples were derived from epithelial surfaces of the ventral part of the trachea). The average AO was found to be deflected in clockwise direction from the TLA by 23° ± 6°. The average difference between the AO and the BBO was determined to be 19° ± 2°. The fact that the direction indicated by the basal foot does not coincide with the axonemal orientation clearly disproves the so far commonly accepted presumption that the effective stroke direction can equivalently be inferred from the ciliary ultrastructure by either the AO, or the BBO.

In order to reinterpret the meaning of these two structurally derived angular observables for mucociliary function, we related them to functional observables, i.e. to the mucociliary transport direction and to the wave propagation direction, which were previously determined on bovine trachea explants by using reflection contrast microscopy Burn (2009). The microscopy setup as well as the applied image processing methods were previously described in Ryser et al. (2007) and subsequently employed in Schätz et al. (2013); Wyss et al. (2018). Relating the structural orientations AO and BBO to the three characteristic functional directions: 1) the sampling-corrected circular mean mucociliary transport direction of 16.1° ± 5.4°, 2) the sampling-corrected circular mean direction of the mean of harmonic plane waves propagating into the laryngeal direction, which was found to be 16.7° ± 6.3°, and 3) the sampling-corrected circular mean propagation direction of the mean wave of 11.7° ± 6.8° it becomes evident that only the AO indicates the effective stroke direction, which is in line with the mucociliary transport and the direction of the harmonic mean plane wave propagation into the laryngeal direction.

The interpretation of the direction of the harmonic plane wave propagating into the laryngeal direction is however highly complex, as the muocociliary wave field exhibits its peculiarities. It must be pointed out that we did not directly measure the direction of the ciliary effective stroke, neither structurally nor functionally. But, in consideration of the noticeable agreement between the propagation direction of the proportion of harmonic plane waves propagating towards the larynx, the mucociliary transport direction and the axonemal orientation, it is highly probable that the axonemal orientation indeed corresponds to the effective stroke direction, which consequently coincides with the transport direction. In order to reach a definitive statement about the accuracy of the coincidence between the axonemal orientation and the effective stroke direction, future functional studies might extract the direction of the effective stroke from, for instance, the optical flow Quinn et al. (2015) in recordings taken by reflection contrast microscopy.

The difference found between the orientation of the basal foot (BBO) and the mucociliary transport direction of 16.1° ± 5.4° makes it very unlikely that the basal foot indicates the direction of transport. This however raises new questions about the orientation of the basal foot: why is it almost perfectly aligned with the TLA? If it is brought about by the external flow, why did we find a considerable disagreement with the transport direction? It is conceivable that the basal foot appendage is structurally related, in a fixed manner, with the axonemal orientation. On the other hand, the beating pattern of individual cilia are generally thought to result from interactions with the environmental fluid. Therefore, the angular deviation between the AO and the BBO could be related in an adaptive manner, as the AO might result from hydrodynamic interactions, and thus, represent an indicator of the prevailing rheology. In a similar morphofunctional study performed on the ciliated epithelium of the bovine oviduct Schätz et al. (2013), we have recently shown that the AO is exactly aligned along the longitudinal axis of the fallopian tube. Furthermore, compared to the mucociliary epithelial surface of the airways, altered rheological conditions are found in the fallopian tube. It specifically lacks of an air-liquid interface and ciliated cells are submerged in a rather aqueous fluid. Future simultaneous measurements of the ciliary AO and the ciliary BBO in the fallopian tube, might therefore bring clarity to the relation between the alignment of the two ultrastructural features. We suspect that the BBO either amounts to roughly −20° (fixed AO-BBO relation), or points into the same direction as the AO, meaning that the AO adaptively results from the prevailing fluid mechanical interactions.

It is known that the basal foot interacts with the apical cytoskeleton (e.g. Vladar and Axelrod (2008); Herawati et al. (2016); Tateishi et al. (2017); Mirvis et al. (2018)). In Herawati et al. (2016), it was specifically shown that the orientation of the basal foot is correlated with the positioning process of cilia. Furthermore, the basal body alignment was found to be disrupted in basal foot depleted (Odf2) mutant mice, and as the basal body alignment was found to be correlated with its orientation, it is probable that the interaction of the basal foot with the cytoskeletal network is crucial for the positioning and orienting process of cilia.

One of our primary aim was to assess whether the mean CO varies between neighboring cells. We reported the median absolute intercellular difference in CO, which amounts to 5° for AO-as well as BBO-inferred values. In order to provide further information on how intercellular orientational deviations relate to intracellular deviations, pairs of cilia having an identical cellular affiliation were compared to cilia pairs having distinct cellular affiliations. This yielded almost identical values for the overall median angular deviations within pairs drawn from the same cell and pairs drawn from distinct cells: for BBO-inferred values the deviation between medians amounts to 0.6° and 1.6° for AO-inferred values. Moreover, we determined 99%-confidence intervals for the difference between intracellular means of neighboring cells. The difference between intracellular means was not significant (*α* = 1%) for three-fourths of all pairs of cells.

We used spatial analysis methods in order to detect possible spatial regularities in CO. In particular, extensive usage of a first-order version of the variogram, the so-called madogram, revealed an intracellular gradual shift in CO. In approximately 90% of all the cells, this ubiquitous intracellular orientational gradient indicated an angular shift in clockwise direction when seen from the right to the left transversely to the TLA.

The emergence of collective ciliary beat patterns has recently been studied in terms of numerical models representing hydrodynamically interacting cilia arrays (e.g. Elgeti and Gompper (2013); Niedermayer et al. (2008); Guirao and Joanny (2007); Ghorbani and Najafi (2017)). These studies typically investigate the effect of geometrical parameters, such as the orientation and the spatial alignment of cilia. An intracellular gradient in CO is expected to strongly affect the emergence as well as the robustness of collective ciliary beating patterns and trigger metachronal waves. It is important to note that the intracellular orientational gradient might, itself, be the result, as well as the cause, of self-organizing processes during development.

Recently, the alignment process of cilia during cell differentiation of cultured multiciliated murine tracheal cells was studied Herawati et al. (2016). A long-term live-imaging system was used in order to track the position and orientation of green fluorescent protein-centrin2-labeled basal bodies during the differentiation process of ciliated cells. It has been shown that the alignment process undergoes four phases, which have been identified in terms of four stereotypical basal body alignments. Relevant here is that the refinement process of the positions of cilia was correlated with the orientation of basal bodies. As the spatial alignment of cilia, therefore, seems to be interrelated with their orientation, future studies might consider the role of intracellular orientational gradients for ciliary alignment, and vice versa.

The mechanism establishing the herein reported gradual shift in cilia orientation within cells remains to be discovered. We suspect a tight relation to planar cell polarity (PCP) proteins, whose asymmetrical localization has recently been demonstrated Vladar et al. (2012); Guirao et al. (2010), as well as to interactions between PCP components and the actin- and microtubular cytoskeletal network of the apical cellular membrane Spassky and Meunier (2017); Vladar et al. (2012); Werner and Mitchell (2012); Werner et al. (2011); Mitchell et al. (2007).

In summary, this study connects structural and functional observables and reveals that out of the axonemal and the basal body directions only the AO points in the direction of the mucociliary transport which corresponds to the direction of the effective stroke. On the other hand, the BBO was found to be almost parallel to the tracheal long axis. Considering the orientation of the cilia on a single cell, a gradual shift was found, whereas the cilia orientation does not change between neighboring cells. The present study interrelates ultrastructural features of individual cilia and their multiscale spatial alignment to mucociliary function and, thus, bridges the gap between ciliary morphology and function, which sheds new light on the mechanisms governing the spatial organization of ciliary activity.

## Materials & Methods

### Sample Preparation

This study was performed with tracheal epithelia collected from four freshly slaughtered, healthy adult cows. Immediately after slaughtering, the trachea was dissected free, opened longitudinally and rinsed with isotonic phosphate buffered saline (PBS). In order to preserve the information of the tissue orientation (i.e. the longitudinal axis of the trachea), large asymmetric trapezoidal epithelium samples (≈ 3cm×2cm) were excised from the tracheal wall (Fig.1). Subsequently, the tissue samples were immediately fixed with 2.5% glutaraldehyde (Agar Scientific, Stansted, Essex, UK) in 0.1M cacodylate buffer (Merck, Darmstadt, Germany), pH 7.4.

For serial block face scanning electron microscopy, two small trapezoidal samples (≈ 3mm × 2mm), were carefully excised from one large trapezoid: one derived from a region overlying a cartilage ring (cartilago trachealis ≙ CT) and one associated with the adjacent annular ligament (ligamentum anulare ≙ LA), as illustrated in Fig.1. The tissue samples were processed according to a specific protocol for serial block face scanning electron microscopy: first, the tissue was rinsed four times with 0,15M Na-cacodylate. The last rinse was supplemented with 0.5% Triton x-100 (Octoxinol 9 (C_14_H_22_O(C_2_H_4_O)n, Sigma-Aldrich, Buchs, Switzerland), a surfactant allowing for a better permeability of contrast agent into cilia. After another wash with 0,15M Na-cacodylate, the staining agents 2% OsO4 (Osmiumtetraoxid, Electron Microscopy Sciences, Hatfield, PA, USA) and 3% potassium ferrocyanide (C_6_N_6_FeK_4_, Sigma-Aldrich, Buchs, Switzerland) in 0.15M Na-cacodylate were added. The tissue was then rinsed with bidistilled water, and pyrogallol (C_6_H_6_O_3_, Sigma-Aldrich, Buchs, Switzerland) was added to enhance the staining of the membrane. Thereafter, samples were successively incubated in 2% OsO_4_, 1% uranyl acetate (Sigma-Aldrich, Buchs, Switzerland) and Waltonfis lead aspartate (Sigma-Aldrich, Buchs, Switzerland) Walton (1979). Between each of the latter steps, the tissue was rinsed with bidistilled water. After this, the tissue was dehydrated through an ascending ethanol series (20%, 50%, 70%, 90%, and twice 100%). The tissue was infiltrated with Durcupan (epoxy resin, Sigma-Aldrich, Buchs, Switzerland) with decreasing concentrations of ethanol (1:3, 1:1, 3:1, pure Durcupan). The resin was polymerized for 48 hours at 60°C.

### Image Acquisition & Analysis

Care was taken in positioning the samples to ensure that the image vertical axis corresponded to the tracheal long axis (TLA), with the laryngeal end directed towards the image top. Threedimensional (3D) ultrastructural images were produced on a Qanta FEG 250 scanning electron microscope (FEI, Eindhoven, The Netherlands) equipped with a 3View2XP serial block face module (Gatan, Munich, Germany) using the Digital Micrograph program (Gatan). In order to achieve an approximately orthogonal sectioning of the cilia, the cutting plane was chosen with great care in the 3view system and was oriented tangentially to the curved cellular surface. Block face sectioning was done from the tips of cilia towards their rootlets with a step size of 120nm (covering a vertical distance of≈ 12*μ*m). The surface of the sample blocks was imaged by back-scattered electron detection. Each image represents a field of view (FOV) with an area of 12×12 *μ*m^2^, which was imaged with a resolution of8192×8192 pixels^2^.

Thus, we obtained 16 stacks of images. Each stack was derived from a sample block volume of 12 × 12 × 12 *μ*m^3^ comprising about 100 sequential serial sections or ≈ 8192 × 8192 × 100 voxels corresponding to a voxel size of 1.5 × 1.5 × 120 nm^3^. The image sequences were processed with the IMOD software package Kremer et al. (1996). CO measurements were done in 3dmod software (image display and modeling program of IMOD), after relative alignment and denoising by nonlinear anisotropic diffusion Frangakis and Hegerl (2001).

### Determining Ciliary Orientation

We used two angular observables to quantify the orientation of cilia with respect to the TLA: 1) the direction of the axonemal orientation (AO) Satir and Christensen (2007); Schätz et al. (2013); Satir et al. (2014); Satir (2016) and 2) the direction of the basal body orientation (BBO) (e.g. Satir (2016)), both considered to determine the direction of the effective stroke.

#### Axonemal Orientation (AO)

The ciliary beating plane is commonly assumed to be orthogonal to the line connecting the central microtubule pair. The ciliary effective stroke is then assumed to point from the microtubular doublet 1 towards the gap between the microtubular doublets 5 and 6 (Fig.7A). This direction with respect to the tracheal long axis was determined from three coordinates: two labelling the central microtubule pair (point ‘a’ and ‘b’ in Fig.7B) and a third labelling the gap between the microtubule doublets 5 and 6 (point ‘c’ in Fig.7B). Finally, the axonemal orientation (AO) with respect to the TLA was defined as the angle ‘Θ’, which is enclosed by the arrow ‘N’ and the vertical arrow ‘V’ being parallel to the TLA and pointing towards the larynx (Fig.7D).

We found a considerable axonemal twist along the ciliary long axis (as illustrated in Figure 7 – Figure Supplement 1). Its effect on our orientational measurements was minimized by pursuing each single cilium along its longitudinal axis (*z*-axis), in order to determine the axonemal as well as the basal body orientation as close as possible to the apical cell surface.

#### Basal Body Orientation (BBO)

Due to the lack of a quantitative study proving the equivalence of the two orientational observables, we decided to measure the orientation of each cilium two-fold (whenever possible), by means of the AO as well as the BBO. The orientation of the basal foot with respect to the TLA was determined from two coordinates: the first indicating the center of the basal body and the second indicating the tip of the basal foot appendage. As illustrated in Fig.8, the angle *ω*, which is enclosed by the arrow ‘P’ (pointing from the center of the basal body towards the tip of the basal foot) and the vertical arrow ‘V’, represents an unambiguous observable for the basal body orientation.

Finally, the axonemal orientation (Θ) as well as the basal body orientation (*ω*) was reported in terms of the angular deviation from the TLA (pointing into laryngeal direction) according to the convention illustrated in Fig.8B.

### Directional/Circular Statistics

Circular statistics represent a subfield of statistics dealing with data distributed on angular or circular scales. Since we are basically dealing with angles herein, it is important to note that angles are distributed on a circular scale and, thus, the designation of zero, as well as of low and high values, is arbitrary Berens (2009). As a consequence, statistical measures designed for data, which are distributed on a linear scale, can principally not be applied to data distributed on a circular scale.

#### Circular Mean Value

In order to calculate the circular mean 〈*γ*〉_*c*_ of a set of *N* angles: (*γ*_1_, *γ*_2_,…, *γ_N_*), each angle *γ_i_* was first transformed into the unit vector 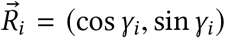. The mean resultant vector 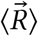 Berens (2009) is then given as

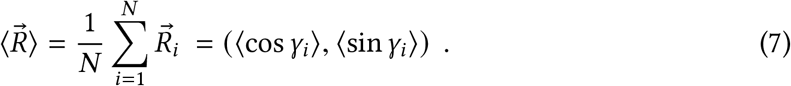

The direction 〈*γ*〉_*c*_ of the mean resultant vector 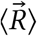, finally represents the mean direction:

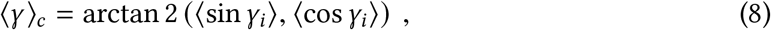

where arctan2(*x*) denotes the 2-argument arctangent function, which is commonly used to calculate the unique inverse of the tangent, as it takes care of the necessary quadrant correction. Note that in Eq.7 and Eq.8, as well as in the following, ‘linear averaging’ is denoted by 〈…〉, whereas circular averaging is denoted by 〈…〉_*c*_.

#### Circular Standard Deviation

The circular analogue to the ‘linear’ standard deviation can be derived from the length of the mean resultant vector 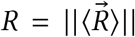, which lies within the interval [0, 1] and represents a measure for circular spread: the narrower the sample {*γ_i_*}_*i*∈{0,1,…,*N*}_ is distributed around its mean direction, the closer *R* is to 1. As the variance is a measure for spread of values in a dataset, the circular variance *V_c_* can be defined according to Berens (2009): *V_c_* = 1 − *R*. The circular standard deviation is commonly defined as Berens (2009)

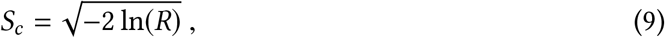

and corresponds to the standard deviation of the wrapped normal distribution.

### Experimental Procedure and Imaging Setup for the Functional Analysis

Fresh tracheas were obtained from the local slaughterhouse. In order to keep track of the tracheal orientation, the tracheas were removed together with the larynx. The tracheas were sealed in plastic bags and transported at temperatures below 10°C in cooling boxes. Upon arrival at the university, they were kept in the refrigerator until the measurements were performed (within approximately 2 hours after the slaughter). Tracheal pieces of sizes between 2 × 2 cm^2^ and 5 × 5 cm^2^ were cut out of the trachea and mounted in a customized sample holder, which was then adequately oriented and aligned on the microscope stage. The microscope stage was equipped with an in-house constructed climate chamber, which allowed to take measurements at controlled environmental conditions. The temperature and air moisture were kept constant and set to 30°C and beyond 90% relative humidity, respectively. The observation of the oscillating mucous surface relied on an epidark field reflectometry setup built around an upright microscope (Nikon Eclipse E600FN, Kanagawa, Japan). Structured illumination (LED illumination delivering an annular hollow beam) through the objective provided adequate contrast while avoiding the direction artifacts associated with the traditional differential interference contrast (DIC). Recordings were taken with the digital CCD camera Dalsa CA-D1T at an image depth of 12 bits at a frame rate of 500 Hertz. Finally, an area of approximately 140 × 140 *μ*m^2^ was captured using a 40×-objective (Nikon Plan Fluor, NA 0.6, WD 2.7 mm).

### Characterization of Observed Wave Fields and Mucociliary Transport

In order to quantitatively characterize the observed mucociliary wave fields, we used sophisticated image processing methods, which were presented in great detail in Ryser et al. (2007) and subsequently used for the analysis of mucociliary phenomena on explants of the bovine fallopian tubes Schätz et al. (2013), as well as for the investigation of the mucociliary clearance on trachea explants of diseased snakes Wyss et al. (2018). In the following the two morphological measurements of the ciliary orientation are compared to the direction of transport as well as to the wave propagation direction. The transport velocity was measured by tracking applied tracer particles (mushroom spores). On each captured image sequence, several particles were tracked. The mean particle velocity (measured in [*μ*m/s]) provides an adequate estimate for the flow velocity, which we denote as 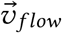 in the following.

The wave propagation velocity was determined two-fold: 1) by the exploration of the space-time correlations contained in each image sequence and 2) by decomposing the image sequences as a synthesis of harmonic plane waves.

The first method delivers an estimate for the mean propagation of the captured surface wavelets 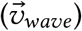, by actually pursuing the temporal displacement of the mean dynamic structure. This was done by determining the shifting velocity of the peak in spatio-temporal cross-correlograms. The second method is based on the generation of the spatial power spectral density, i.e. the distribution of the wave vectors 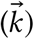, from which we determined the mean wave vector propagating into the pharyngeal direction 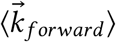 and the mean wave vector propagating into the opposite direction 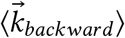.

## Acknowledgments

We would like to thank our former collaborators, Manuel Ryser and Andreas Burn, for the research they performed on mucociliary transport. This work was supported by the Swiss National Science Foundation (grant #163761 to B.Z.). Microscopy was performed on equipment supported by the Microscopy Imaging Center (MIC), University of Bern, Switzerland.

## Author Contributions

Martin Schneiter: conceptualization, data curation, formal analysis, investigation, methodology, software, writing - original draft and writing - review & editing; Sebastian Halm: conceptualization, data curation, investigation, methodology, resources and writing - original draft; Adolfo Odriozola: data curation, investigation, methodology and resources; Helga Mogel: investigation, methodology and resources; Jaroslav Rička: conceptualization, methodology and supervision; Michael H. Stoffel: conceptualization, investigation, methodology, project administration, resources, supervision and writing - review & editing; Benoît Zuber: conceptualization, data curation, funding acquisition, investigation, methodology, project administration, resources, supervision and writing - review & editing; Martin Frenz: conceptualization, investigation, methodology, project administration, resources, supervision and writing - review & editing; Stefan A. Tschanz: conceptualization, data curation, investigation, methodology, project administration, supervision, writing - review & editing

---

**Table 1 – Figure Supplement 1:**
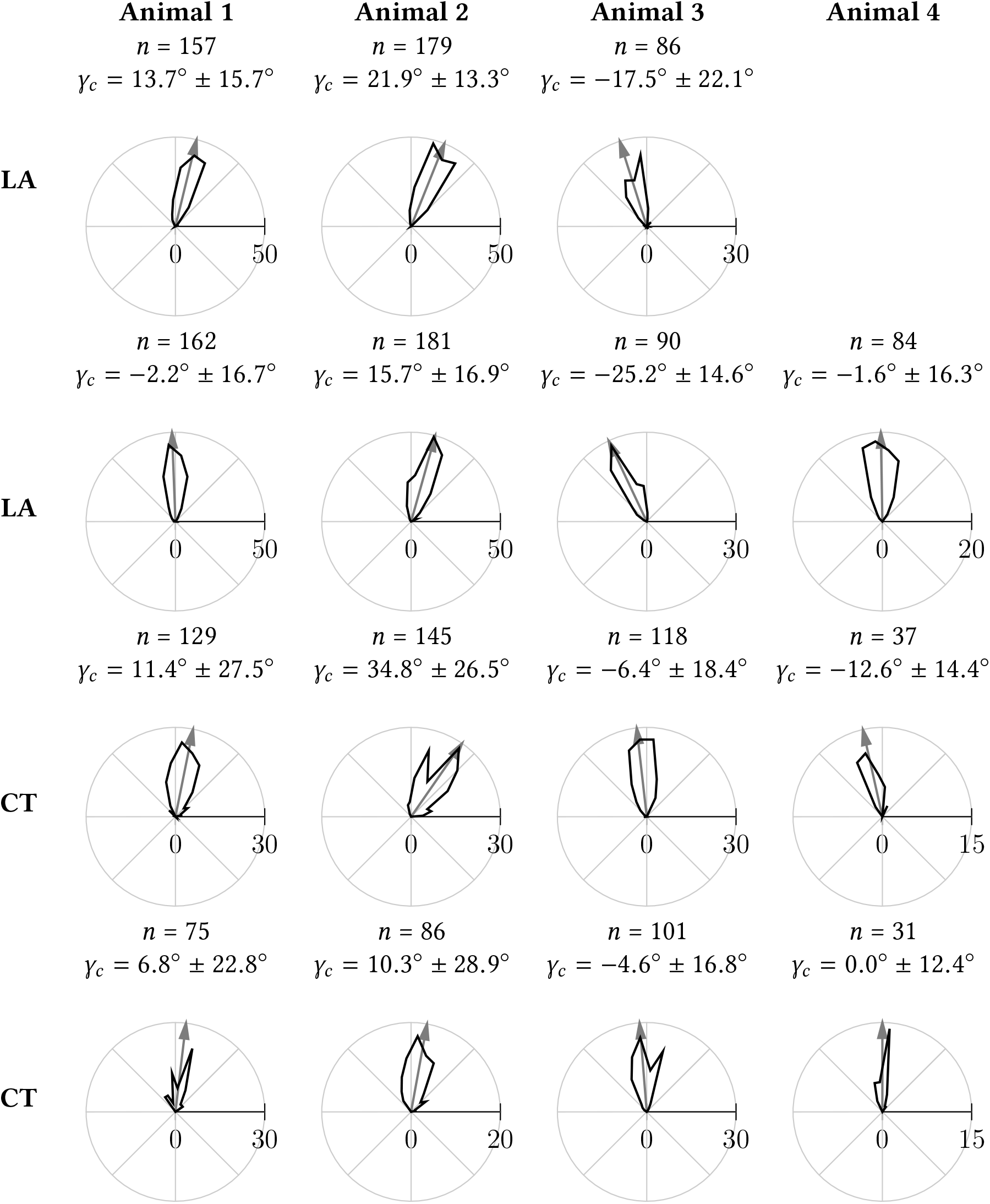
Each polar histogram (black curve) illustrates the distribution of the ciliary power stroke directions as inferred from the basal body orientation in a field of view. The arrow indicates the circular mean direction. The histograms were created by using a bin width of 10°.

**Table 2 – Figure Supplement 1:**
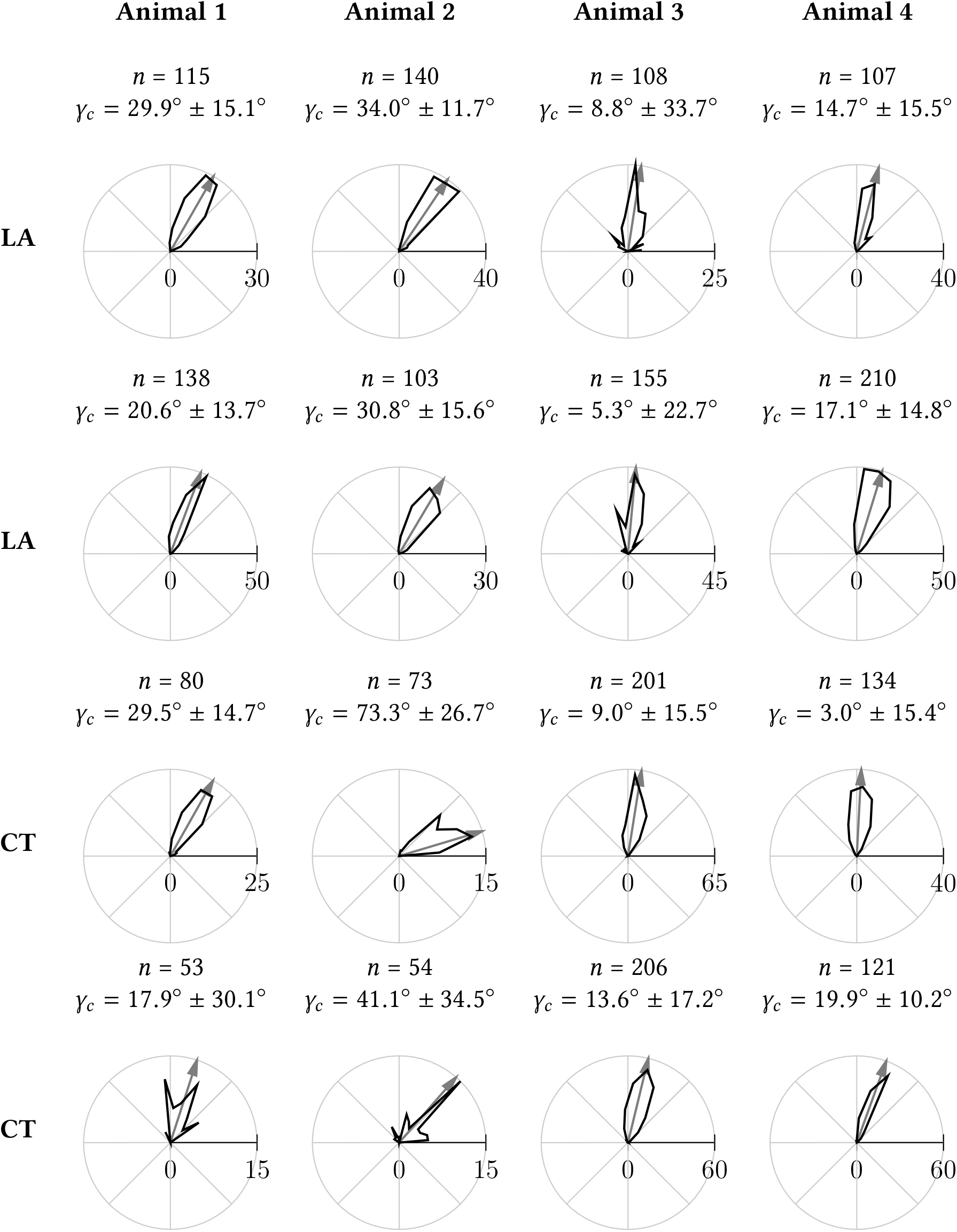
Each polar histogram (black curve) illustrates the distribution of the ciliary power stroke directions as inferred from the axonemal orientation in a field of view. The arrow indicates the circular mean direction. The histograms were created by using a bin width of 10°.

**Figure 6 – Figure Supplement 1:**
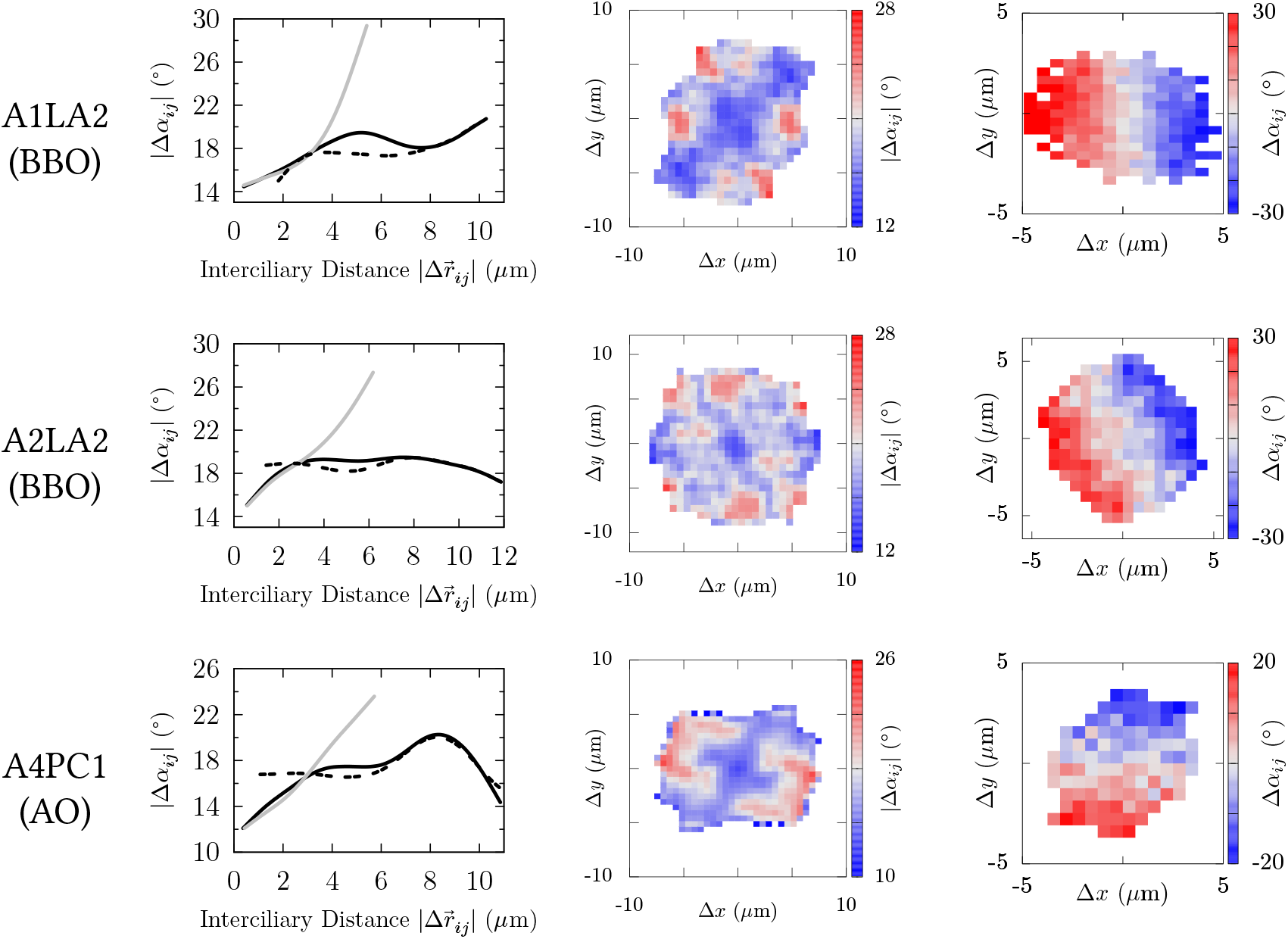
Left panels: radial variograms (intra-FOV: solid black, intracellular: solid gray, intercellular: dashed black). Middle panels: absolute directional FOV-variograms. Right panels: signed directional intracellular variograms.

**Figure 7 – Figure Supplement 1:**
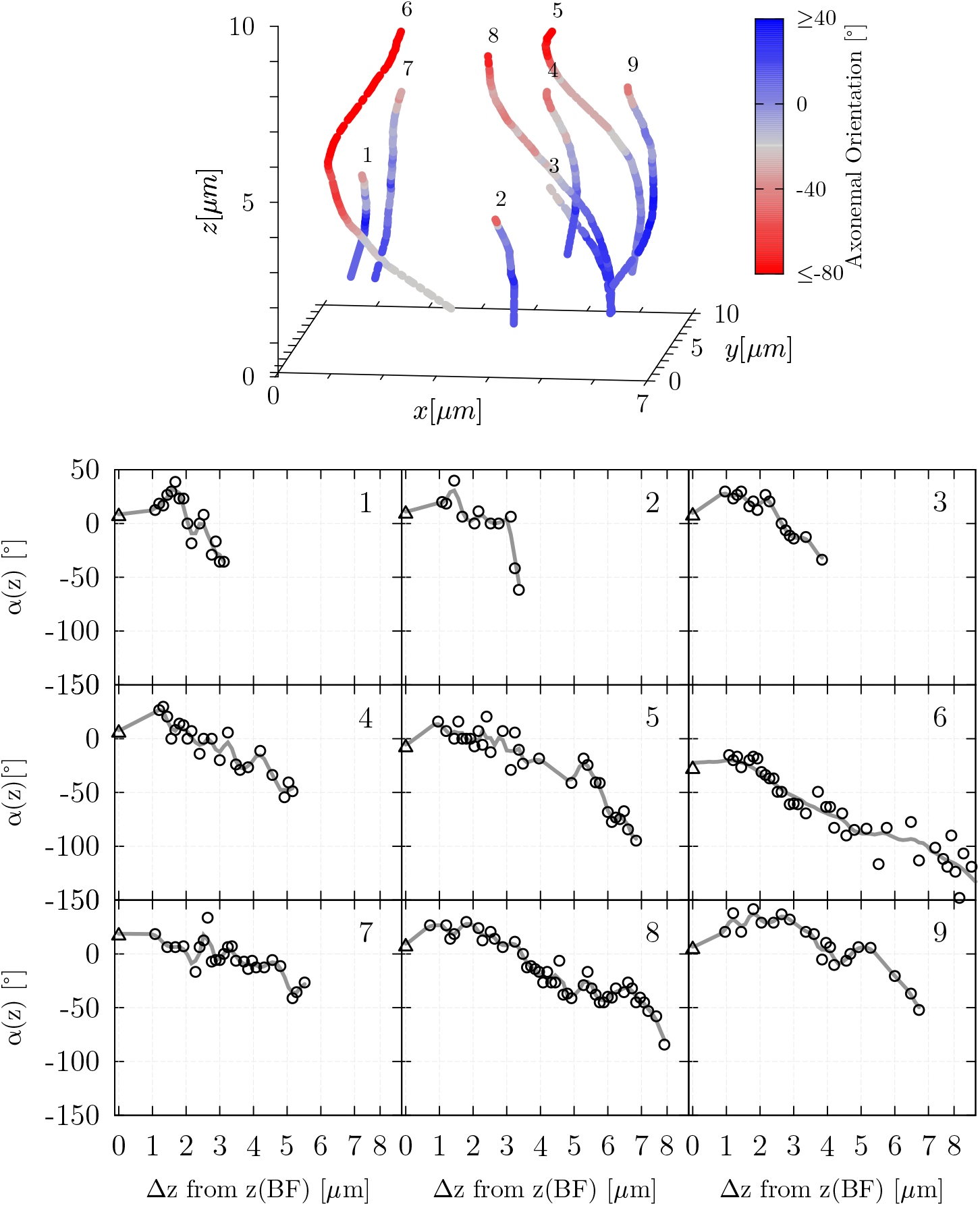
The graphs illustrate the variation of the effective stroke direction along the ciliary long axis (z-axis) for nine cilia derived from the same sample (A1LA2). The (quasi)three-dimensionalgraphatthe top illustrates the (right-handed)ciliarytwistalong the ciliary long axis. The lower graph presents the angular measurements of cilium 1-9 illustrated at the top (triangles correspond to BBO-inferred values and circles to AO-inferred values). The abscissa indicates the vertical distance from the cutting plane in which the basal foot orientation was measured. The grey line results from interpolating and slightly smoothing the data sets. As it can be seen, the effective stroke direction as inferred from the axonemal orientation varies considerably along the ciliary long axis. Therefore, great care was especially taken in order to measure the axonemal orientation as close as possible to the apical cell surface.

